# Cancer cells adapt FAM134B-BiP complex mediated ER-phagy to survive hypoxic stress

**DOI:** 10.1101/2021.02.05.429931

**Authors:** Sandhya Chipurupalli, Raja Ganesan, Giulia Martini, Luigi Mele, Elango Kannan, Vigneshwaran Namasivayam, Vincenzo Desiderio, Nirmal Robinson

## Abstract

In a tumor microenvironment cancer cells experience hypoxia resulting in the accumulation of misfolded/unfolded proteins in the endoplasmic reticulum (ER) which elicit unfolded protein response (UPR) as an adaptive mechanism. UPR activates autophagy enabling the degradation of misfolded/unfolded proteins. More recently, ER-specific autophagy has been implicated in the removal of damaged ER and restoration of ER-homeostasis. Our investigations reveal that during hypoxia induced ER-stress, the ER-phagy receptor FAM134B targets damaged portions of ER into autophagosomes to restore ER-homeostasis in cancer cells. Loss of FAM134B in breast cancer cells results in increased ER-stress and reduced cell proliferation. Mechanistically, upon sensing hypoxia activated proteotoxic stress, the ER chaperone BiP forms a complex with FAM134B and promotes ER-phagy. Our studies have further led to the identification of a pharmacological agent vitexin that disrupts FAM134B-BiP complex thereby inhibits ER-phagy and suppresses breast cancer progression in vivo.

## Introduction

Cancers often encounter a characteristic microenvironment called tumor microenvironment (TME) which comprises of a chemical microenvironment (pH, hypoxia, metabolite concentration) and a cellular microenvironment (blood vessels, immune suppressor cells, fibroblasts, extracellular matrix, stromal cells) which influences the growth of cancerous cells ^1, 2, 3, 4^. Hypoxic environment arises as a result of vascular insufficiency during the tumor expansion and progression^5^. It alters the cancer cell metabolism and contributes to therapy resistance by activating certain adaptive responses such as endoplasmic reticulum (ER) stress, anti-oxidative responses and autophagy^6^. Therefore hypoxia is considered as a major impediment for effective cancer therapy ^5, 7^.

ER is a multifunctional organelle, but it is central for protein synthesis, modifications and transport. The disulphide bonds that are formed during protein synthesis are independent of oxygen availability whereas, the bonds that are formed during the post-translational folding in the ER are oxygen-dependent^8^. This process is altered during hypoxia resulting in the accumulation of misfolded/unfolded proteins in the ER, perturbing its homeostasis. Thus, hypoxia directly impacts protein modifications in the ER resulting in the activation of unfolded protein response (UPR) to preserve ER homeostasis^9^. UPR is a signaling system which activates cellular responses coordinated via three key regulators – inositol-requiring enzyme 1 (IRE1), PKR-like ER kinase (PERK) and activating transcription factor 6 (ATF6)^10, 11, 12, 13^. Binding immunoglobulin protein (BiP or glucose-regulatory protein 78 – Grp78) is a chaperone present abundantly in the ER which transiently binds to the luminal domain of UPR receptors – IRE1, PERK and ATF6^14^. When the misfolded/unfolded proteins begin to accumulate in the ER, BiP rapidly dissociates from the three UPR signaling sensors and binds the exposed hydrophobic regions of the nascent polypeptides to facilitate proper folding. In addition, UPR also induces autophagy as a key response to the stress pathway activation in cancer cells which allows them to maintain metabolic homeostasis^15, 16, 17^.

Autophagy involves the sequestration of cytoplasmic components into autophagosomes which then fuse with lysosomes and degrade their contents^18^. Although, autophagy is a constitutive homeostatic mechanism which regulates intracellular recycling, it is also a major stress responsive mechanism that facilitates the removal of damaged proteins and organelles^19^. Thus, autophagy bestows tolerance to stress and sustains cell viability under hostile conditions and is considered a “double-edged sword” because of its ability to suppress tumor yet promote tumor survival under stress^19^. Despite accumulating evidences suggesting that autophagy is critical in cancer, it is still a question of intense debate and remains complex^20^. For a long time autophagy was considered a non-selective degradation pathway however, recent research has revealed that autophagy can selectively degrade specific organelles including mitochondria (mitophagy), peroxisomes (pexophagy), ER (ER-phagy or reticulophagy), nucleus (nucleophagy)^18^ and aggregate-prone proteins (aggrephagy)^21^.

ER-phagy was first described by Peter Walter’s group where they demonstrated that selective engulfment of ER into autophagosomes utilize several autophagy proteins induced by UPR and these are essential for the survival of cells exposed to severe ER stress ^22^. ER was originally considered as the primary source of autophagosome membranes ^23^ and ER membranes observed in autophagosomes was viewed as a result of bulk engulfment of cytosol ^23, 24^. Lately, identification and characterization of receptors that mediate the specific elimination of ER through autophagosomes have been identified and termed “ER-phagy or reticulophagy” ^25^. To date, seven ER-resident proteins have been identified as selective ER-phagy receptors: FAM134B ^26^, RTN3 ^27^, SEC62 ^28^, CCPG1 ^29^, ATL3^30^, TEX264 ^31, 32, 33^ and the recently identified soluble ER-phagy receptor CALCOCO1^34^. FAM134B and RTN3 are shown to have a crucial role in degrading the ER in response to severe stress or starvation. Mutations in the genes coding for the ER-phagy receptors have been associated with various pathologies but the process of ER-phagy has not been implicated in pathologies till date^35^.

Here we report that hypoxia induced ER-stress in cancer cells activates ER-phagy and is specifically orchestrated by the ER-phagy receptor FAM134B. Upon hypoxic stress FAM134B complexes with the ER chaperone BiP to target ER for autophagic degradation. Inhibition of ER-phagy by silencing FAM134B or BiP reduces breast cancer cell proliferation. Furthermore, we have identified vitexin as a pharmacological inhibitor of FAM134B-BiP complex regulated ER-phagy which is also able to reduce the tumor burden in a breast cancer xenograft model.

## Results

### Hypoxia induces ER stress response and autophagy

Hypoxia is characterized by the stabilization of HIF-1a hence, we first investigated if HIF-1a is expressed and stabilized in our model of MCF-7 breast cancer cells upon hypoxic stress induced chemically using cobalt chloride (CoCl_2_) or growing cells in hypoxic environment (1% O_2_) for 24h. As expected, both chemical induction (CI) **(Fig 1a)** and hypoxic environment (HE) **(Fig 1b)** resulted in HIF-1a expression and stabilization. Although CoCl_2_ has been widely used as a chemical inducer of hypoxia, reports indicate that CoCl_2_ activates a complex relationship between adaptive and cell death responses^36^. We observed HIF-1a expression and cell proliferation were concentration dependent for CI-hypoxia **(Ext Fig 1a)**. Time lapse imaging of MCF-7 cells treated with CoCl_2_ at 500µM for 24h showed more than 2-fold increase in cell proliferation compared to the untreated cells (normoxic cells) **(Fig 1c and movies of normoxic and CI-hypoxic cells in supplementary videos 1 & 2 respectively)**. 500µM of CoCl_2_ was chosen as an optimal concentration which induced HIF-1a and also increased cell proliferation compared to untreated cells for the time course of our experiments.

**Fig 1:**
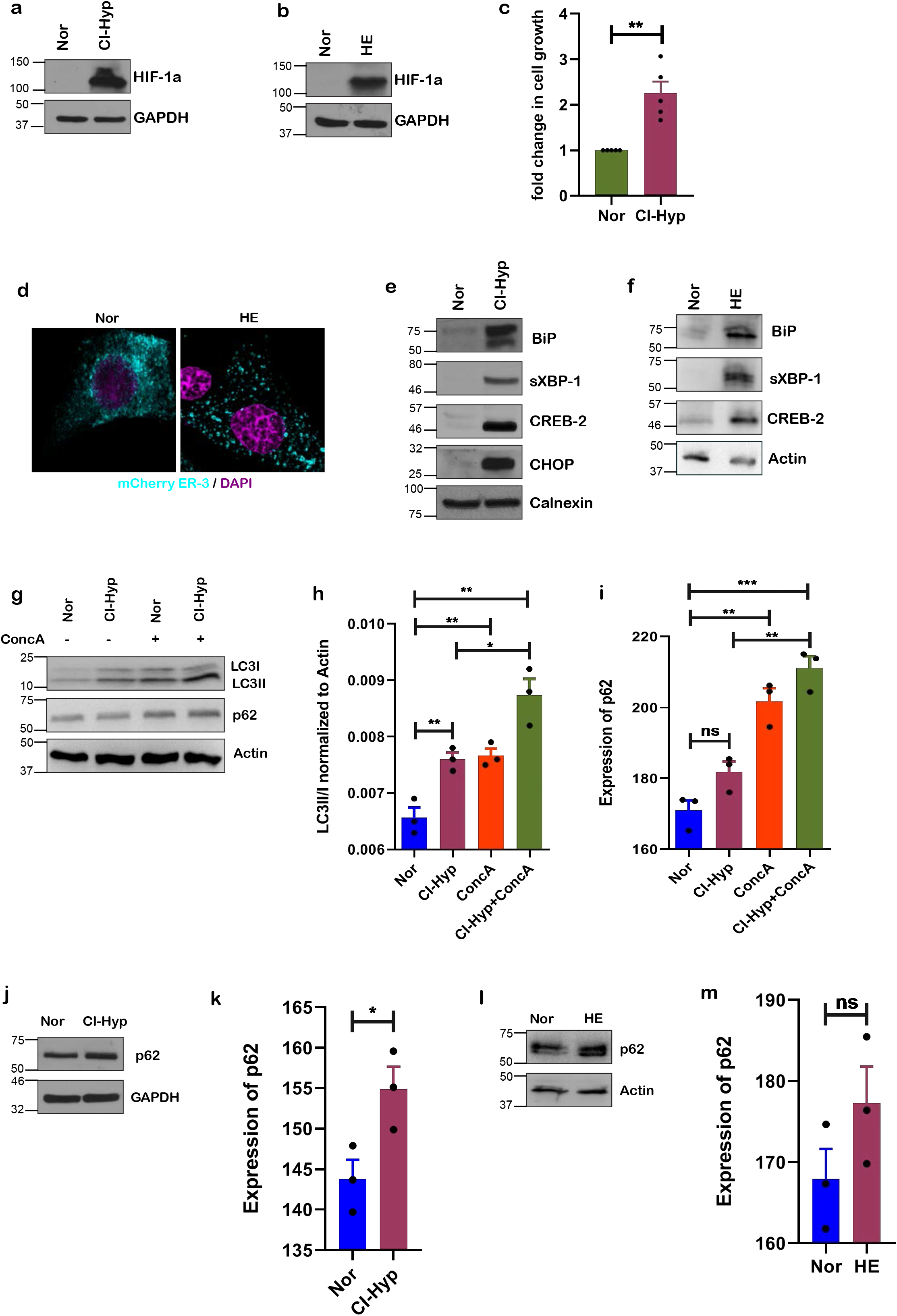
Hypoxia induces ER stress response and autophagy. **(a)** Immunoblot showing stabilization of HIF-1a during CI-hypoxia and **(b)** HE (1% O_2_) *(n=3)*. **(c)** Proliferation of MCF-7 cells upon hypoxia compared to normoxia*(n=5)*. **(d)** Microscopy of MCF-7 cells transfected with mCherry-ER-3 plasmid (Cyan). **(e)** Expression of UPR proteins upon CI-hypoxia and **(f)** cells grown in hypoxic environment (HE) (1% O_2_) compared to normoxia *(n=3)*. **(g)** Immunoblot of LC3I to LC3II conversion and p62 upon concanamycin A treatment during CI-hypoxia in comparison with control *(n=3)*. **(h)** Densitomrtric quantification of LC3I to LC3II conversion and **(i)** p62 upon concanamycin A treatment during CI-hypoxia in comparison to control *(n=3)*. (**j)** Immunoblot of p62 upon CI-hypoxia and **(k)** densitomrtric quantification of p62 upon CI-hypoxia *(n=3)*. (**l)** Immunoblot of p62 in cells grown in HE (1% O_2_) and (**m)** densitomrtric quantification of p62 upon CI-hypoxia and HE (1% O2) *(n=3)*.

Hypoxia results in the accumulation of unfolded/misfolded proteins in the endoplasmic reticulum (ER) causing ER-stress and cancer cells adapt by activating UPR mechanisms which enable them to survive and proliferate^4, 9^. Confocal microscopy of cells expressing mCherry-ER-3 (calreticulin + KDEL) grown in HE showed altered ER structure compared to cells grown in normoxia **(Fig 1d)**. Consistently, quantitative real-time PCR (qRT-PCR) **(Ext Fig 1b-1e)** and Western blot analysis of UPR markers BiP, spliced XBP-1 (XBP-1s), CREB-2/ATF4 and CHOP showed significant upregulation upon CI-hypoxia **(Fig 1e)**. Increased expression of UPR target proteins was also confirmed in cells grown in HE **(Fig 1f)**. These data suggest that hypoxia induced HIF-1a and ER-stress response correlates with increased cancer cell proliferation.

It is well-known that autophagy is required for the survival of hypoxic cancer cells^37^ and it assists in degrading misfolded/unfolded proteins to reestablish ER homeostasis ^4, 38^. Hence, we next investigated whether autophagy is induced in cancer cells under hypoxia. In cells subjected to CI-hypoxia **(Ext Fig 1f and 1h)** and HE **(Ext Fig 1g and 1i)**, LC3B, a *bonafide* marker of autophagy activation, showed increased conversion to its active form (LC3II) compared to the normoxic cells. Moreover, inhibition of lysosomal activity using concanamycin A lead to further increase in the accumulation of LC3II in cells treated with CoCl_2_ **(Fig 1g-1h)**. This was congruent with confocal microscopic analysis of hypoxic cancer cells stained for LC3B which showed increased distribution of LC3B puncta **(Ext Fig 1j and 1k)** compared to normoxic cells. Similarly, presence of LC3B puncta was also observed in MMTV-pyMT mouse breast cancer tissue sections **(Ext Fig 1l)**. Time lapse imaging of MCF-7 cells expressing GFP-tagged-WIPI-1 (GFP-WIPI-1): another autophagosome-marker over a period of 24h showed increased turnover of numerous WIPI-1 positive autophagosomes upon CI-hypoxia **(movies of normoxic and CI-hypoxic cells in supplementary videos 3&4 respectively)**. We also questioned the fate of p62/SQSTM1/Sequestosome-1 an autophagy receptor of macroautophagy which directly binds to LC3B and the cargo and subsequently degrades in the lysosomes^39^. Interestingly, neither CI-hypoxia **(Fig 1j and 1k)** nor HE-hypoxia **(Fig1l and 1m)** altered p62 levels. Concanamycin A treatment led to further accumulation of p62 in both the normoxic and hypoxic cells **(Fig 1g and 1i)**. Taken together, these data suggest that cancer cells survive hypoxic stress by inducing a non-canonical autophagy independent of p62 degradation.

### Hypoxia induces ER-phagy to maintain ER homeostasis

Various types of selective autophagy that are independent of p62 degradation have been described. In these processes, damaged organelles are selectively targeted into autophagosomes for degradation. Since we observed that the ER is stressed during hypoxia **(Fig 1d)**, we questioned if the more-recently described ER-selective autophagy (ER-phagy)^40,41^ was involved in the removal of damaged ER. Microscopic analysis of cells expressing GFP-WIPI-1 and mCherry-ER-3, showed increase in GFP-WIPI-1 puncta (autophagosomes) in cells subjected to hypoxia and they were found to co-localize with mCherry-ER-3 **(Fig 2a & individual channels in Ext Fig 2a)**. Consistent results were obtained from time lapse imaging of CI-hypoxic cells **(representative frames of time lapse imaging at different time points in Fig 2b and the movies of normoxic and CI-hypoxic cells in supplementary videos 5&6)**. Co-localization of WIPI-1 with the ER suggested that damaged ER is engulfed by autophagosomes to mitigate ER-stress caused by hypoxia. However, during macroautophagy ER also provides membrane for autophagosome formation^23, 24^. Hence, we next investigated the expression of ER-phagy-specific receptors such as FAM134B, RTN3, Sec62, CCPG1 and the COPII subunit Sec24C which have been shown to target ER for autophagosomal degradation. We did not observe any change in the expression of CCPG1, RTN3 and Sec24C but a modest increase in Sec62 **(Ext Fig 2b)**. In contrast, FAM134B decreased consistently upon CI-hypoxia and in cells grown in HE **(Fig 2c-2d)**. However, removal of hypoxic stress by replacing CoCl_2_ with medium without CoCl_2,_ restored the expression of FAM134B **(Fig 2e and Ext Fig 2c)**. We found that the decline in FAM134B was due to lysosomal degradation as inhibition of lysosomal activity using concanamycin A prevented the degradation during HE hypoxia **(Fig 2f and Ext Fig 2d)** and CI-hypoxia **(Ext Fig 2e)**. We also observed a similar degradation of FAM134B in U251 glioblastoma **(Ext Fig 2f)** and C32 melanoma **(Ext Fig 2g)** indicating ER-phagy as a general mechanism in cancer cells to counteract hypoxia induced stress. Furthermore, LC3B co-localized with FAM134B in hypoxic cells **(Fig 2g-2i and individual channels in Ext Fig 2h)** and in MMTV-pyMT mouse breast cancer tissue sections **(Fig 2h & individual channels in Ext Fig 2i)**. Moreover, LC3B co-immunoprecipitated with endogenous FAM134B under hypoxic condition while, there was little interaction between FAM134B and LC3B in control cells, strongly supporting an increase in ER-phagy upon hypoxia **(Fig 2j)**. Having found that hypoxia induces ER-phagy, we next examined if FAM134B-dependent ER-phagy contributes to ER-stress and cell proliferation upon hypoxia. As shown previously, CI-hypoxia increased UPR but, knocking down *FAM134B* **(Ext Fig 2j)** further enhanced the induction of UPR proteins **(Fig 2k and Ext Fig 2k-2m)** meaning increased ER-stress. Silencing *FAM134B* modestly reduced cell proliferation under normoxic conditions but was highly reduced under hypoxic conditions **(Fig 2l)**. These observations indicate that hypoxia induces ER-phagy to overcome ER-stress and that FAM134B-dependent ER-phagy is vital for cancer cells to proliferate under hypoxic stress.

**Fig 2:**
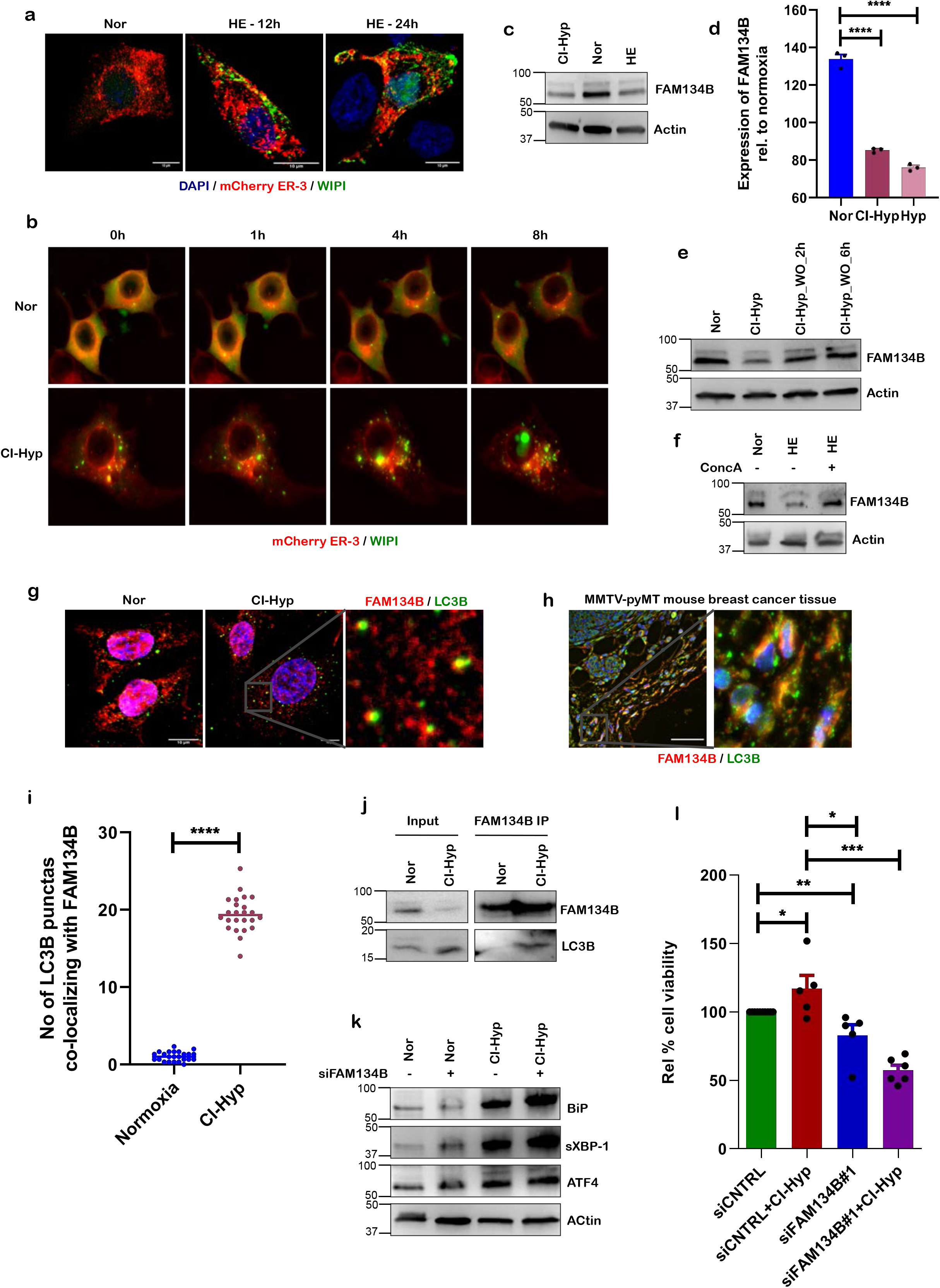
Hypoxia induces ER-phagy to maintain ER homeostasis. **(a)** Confocal microscopical image of MCF-7 cells co-transfected with mCherry-ER-3 & GFP-WIPI plasmid upon hypoxia for 12h and 24h. **(b)** Representative frames from time-lapse imaging of mCherry-ER-3 & GFP-WIPI plasmids co-transfected MCF-7 cells upon CI-hypoxia compared to normoxia *(n=2)*. **(c)** Immunoblot of FAM134B during CI-hypoxia and HE (1% O_2_) compared to normoxia *(n=5)*. **(d)** Densitometric quantification of FAM134B during CI-hypoxia and HE (1% O_2_) compared to normoxia *(n=5)*. **(e)** Western blot analysis of FAM134B expression in the presence of CoCl_2_ and after CoCl_2_ wash out *(n=3)*. **(f)** Western blot of FAM134B during HE (1% O2) in the presence and absence of concanamycin A *(n=4)*. **(g)** Immunofluorescence microscopy of MCF-7 cells stained for FAM134B (red) and LC3B (green) upon hypoxia in comparison with normoxia and **(h)** MMTV-pyMT breast cancer tissue section. **(i)** Quantification of number of LC3 punctas co-localizing with FAM134B *(n=3; punctas were counted from 24 cells in each experiment)*. **(j)** Co-immunoprecipitation of LC3B with FAM134B during hypoxia compared to normoxia *(n=3)*. **(k)** Immunoblot of UPR proteins upon *FAM134B* knockdown using specific siRNAs *(n=3)*. **(l)** Relative % cell viability of hypoxic cells compared to normoxic cells depleted of FAM134B using specific siRNA *(n=5)*.

### Hypoxia induced ER-phagy is BiP dependent

Since we deciphered that hypoxia leads to activation of UPR and FAM134B dependent ER-phagy, we asked if this is HIF-1a dependent. Although depletion of HIF-1a downregulated hypoxia induced UPR **(Fig 3a-3e)**, degradation of FAM134B was not affected in cells subjected to CI-hypoxia **(Fig 3f and 3h)** and HE **(Fig 3g and 3i)**. This suggests that hypoxia-induced ER-phagy is independent of HIF-1a but, triggered by an alternative pathway most likely connected to the presence of misfolded ER proteins. FAM134B lacks an intraluminal domain therefore, we wondered if it senses the accumulation of unfolded/misfolded proteins by interacting with the ER-stress sensor BiP. Indeed, silencing BiP prevented the degradation of FAM134B during hypoxia **(Fig 3j and 3k)** signifying that ER-phagy is BiP-dependent under hypoxic stress. BiP was also found to strongly co-express with FAM134B in MMTV-pyMT mouse breast cancer tissue sections **(Fig 3l and individual channels in Ext Fig 3a)** and in human breast cancer tissues **(Fig 3m and individual channels in Ext Fig 3b)**. Strikingly, BiP co-immunoprecipitated with endogenous FAM134B when MCF7 cells were subjected to hypoxia **(Fig 3n)**. Taken together, these data show that ER-phagy is a specific response to ER-stress and is coregulated by BiP and FAM134B.

**Fig 3:**
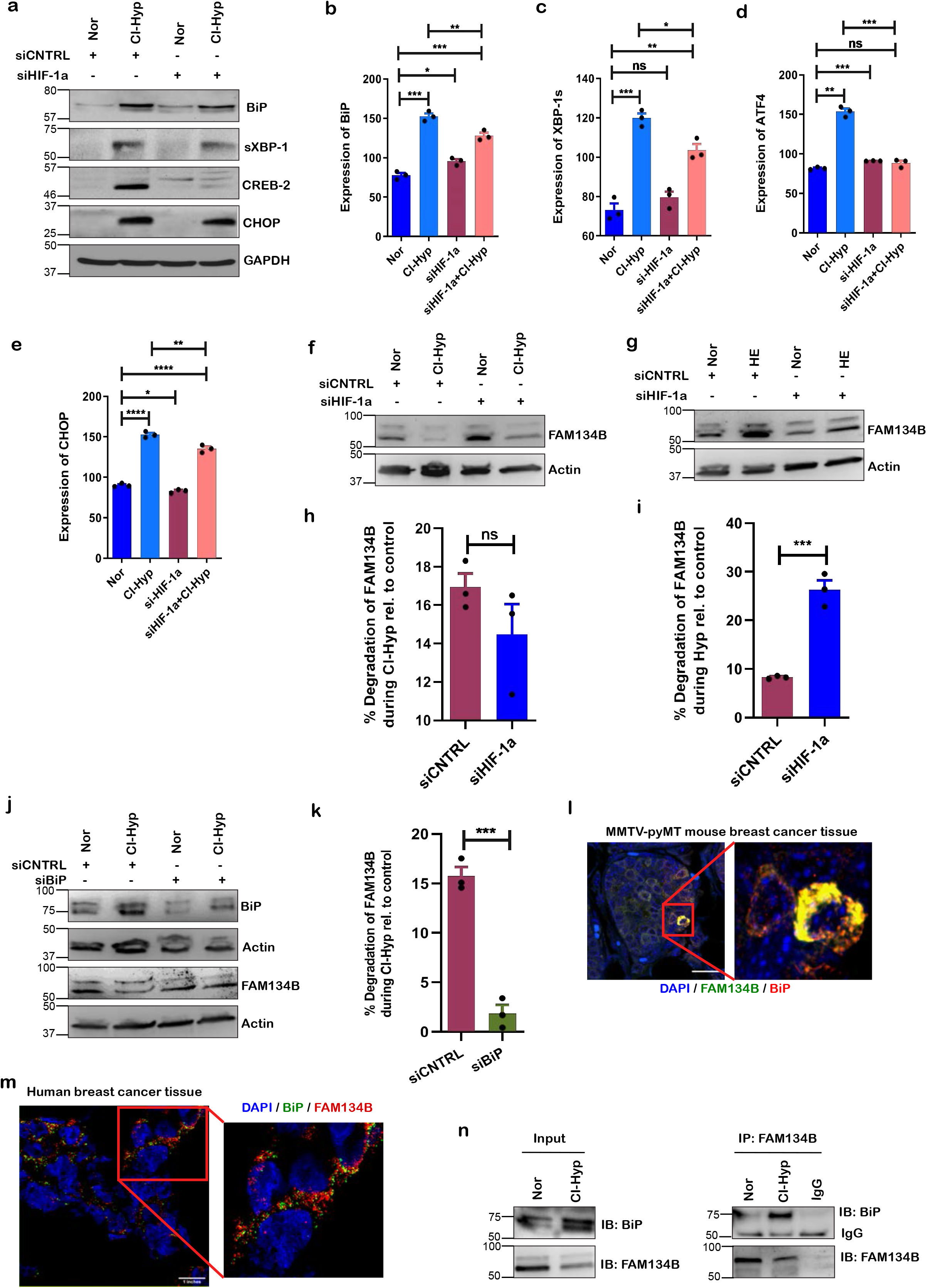
Hypoxia induced ER-phagy is BiP-dependent. **(a)** Immunoblot of UPR proteins in MCF-7 cells transfected with HIF-1a siRNA or control siRNA and subjected to CI-hypoxia *(n=3)*. **(b-e)** Densitometric UPR proteins; **(b)** BiP **(c)** XBP-1s **(d)** ATF4 **(e)** CHOP in MCF-7 cells transfected with HIF-1a siRNA or control siRNA and subjected to CI-hypoxia *(n=3)*. **(f)** Western blot analysis of FAM134B in HIF-1a-depleted cells during CI-hypoxia and **(g)** HE (1% O2) *(n=3)*. **(h)** Densitometric quantification of FAM134B expression in HIF-1a siRNA transfected cells during CI-hypoxia and **(i)** HE (1% O2) *(n=3)*. **(j)** Western blot analysis of FAM134B in BiP knock down cells using siRNA during CI-hypoxia *(n=3)*. **(k)** Densitometric quantification of FAM134B during CI-hypoxia upon silencing BiP *(n=3)*. **(l)** Confocal microscopy of MMTV-pyMT breast cancer tissue section stained for FAM134B and BiP; scale bar = 20uM. **(m)** Immunoblot illustrating co-immunoprecipitation of BiP with FAM134B during hypoxia compared to normoxia *(n=3)*.

### In-Silico molecular docking studies identified vitexin as a potential BiP inhibitor

Above shown results revealed that ER-phagy alleviates ER-Stress response and facilitates the survival and progression of hypoxic cells. Hence, we intended to pharmacologically target BiP mediated ER-phagy to prevent cancer cell proliferation. For this, we have retrieved the high-resolution X-ray crystal structure of the protein from the Protein Data Bank (PDB ID: 5F0X.pdb, Resolution: 1.6 Å). Validation by Ramachandran Plot depicted that 98% and 2% of amino acids are in favorable and allowed regions respectively **(Ext Fig 4a)**. Therefore, we next performed molecular docking studies using Schrödinger Suite 2015-3. A small set of small molecules library was docked onto the binding site of the protein and identified an apigenin flavone glucoside, vitexin as a potential molecule to target BiP. Docking scores are shown in Supplement Table 1. Vitexin showed lowest glide score towards BiP i.e., −8.3Kcal/mol **(Supplement Table 1)**. The putative binding mode of vitexin and important residues in the binding site of the BiP are shown in **Fig. 4a and 4b**. The binding site comprises a higher number of charged and polar residues. Vitexin was bound inside the binding site by two strong hydrogen bonding interactions between OH of the glucoside and the side chain of amino acid residues N389 and R367. The keto group of the flavanone moiety forms hydrogen bond interaction with S300 and arene-hydrogen interaction with R297 **(Fig 4c)**. These predicted interactions of vitexin resulted in top rank based on the docking score meaning higher binding affinity when compared to the other molecules screened against BiP. As a next step, we performed molecular dynamics simulations of the BiP-vitexin complex for explaining the stability of the predicted binding pose in the binding site of the protein and compared with the simulations of the protein structure without the ligand inside the binding site. The calculated root mean square deviation (RMSD) values of the Cα atoms of the complex and the apo structure rapidly reached an equilibrium state with approximately 1 Å deviation from the first frame of 100 ns simulations **(Fig. 4d)**. The visual analysis of the trajectories shows that vitexin was anchored inside the binding site with the interaction pattern identified from the docking studies **(Ext Fig. 4b, Supplementary video Supplementary Video 7)**. It maintains the key hydrogen bond interaction with the three residues (N389, R367 and S300) and possible arene interaction with R297. The root mean square fluctuation (RMSF) value of the protein showed a profile with large fluctuations in the Cα atoms are more stabilized in the BiP in complex with vitexin when compared to the protein without the binding of vitexin in the binding site **(Fig. 4e)**. Based on these findings using molecular docking and molecular dynamics approaches, we subjected vitexin for further in-vitro validation.

**Table 1:**
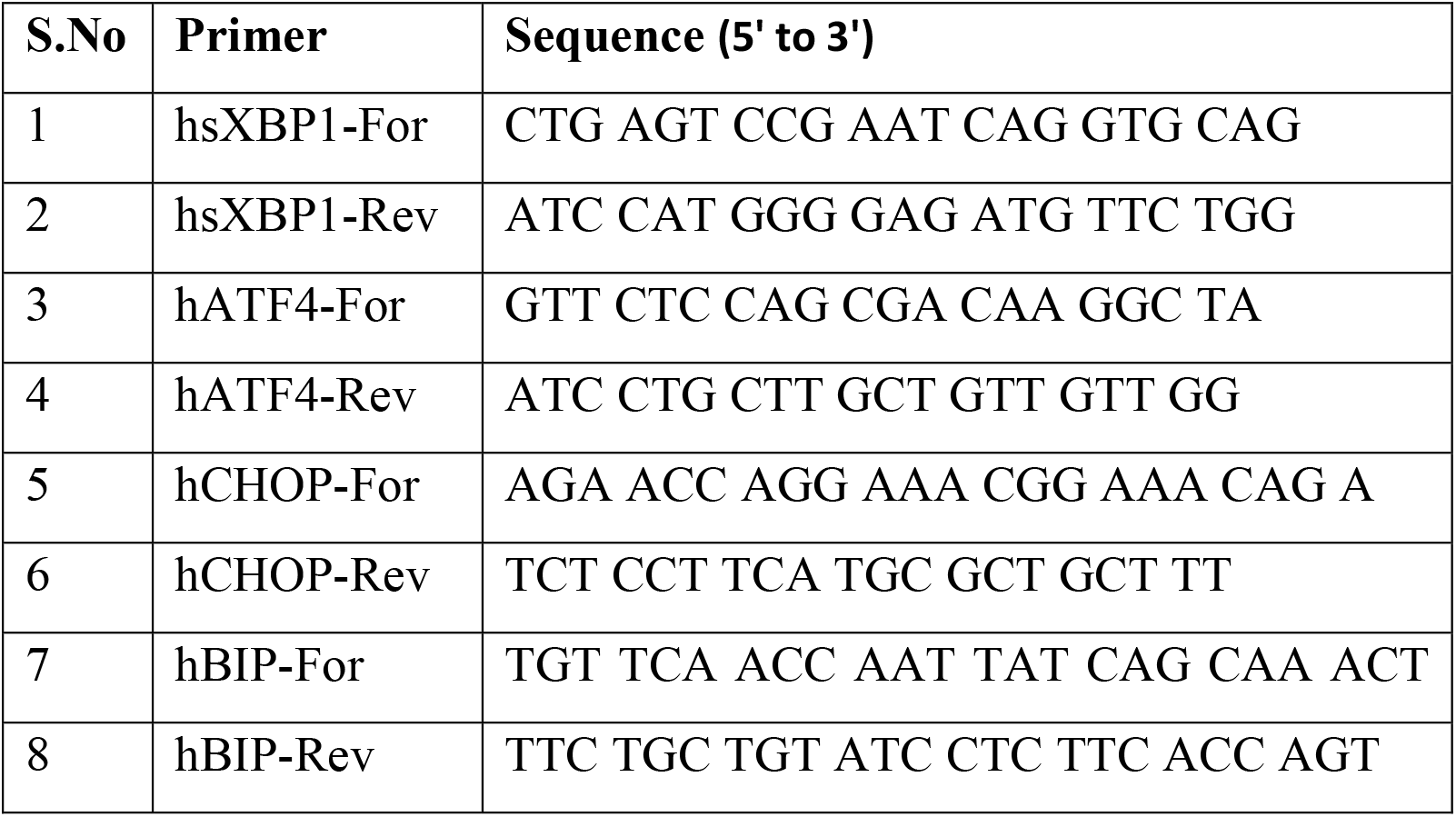
Primers used in the study

**Fig 4:**
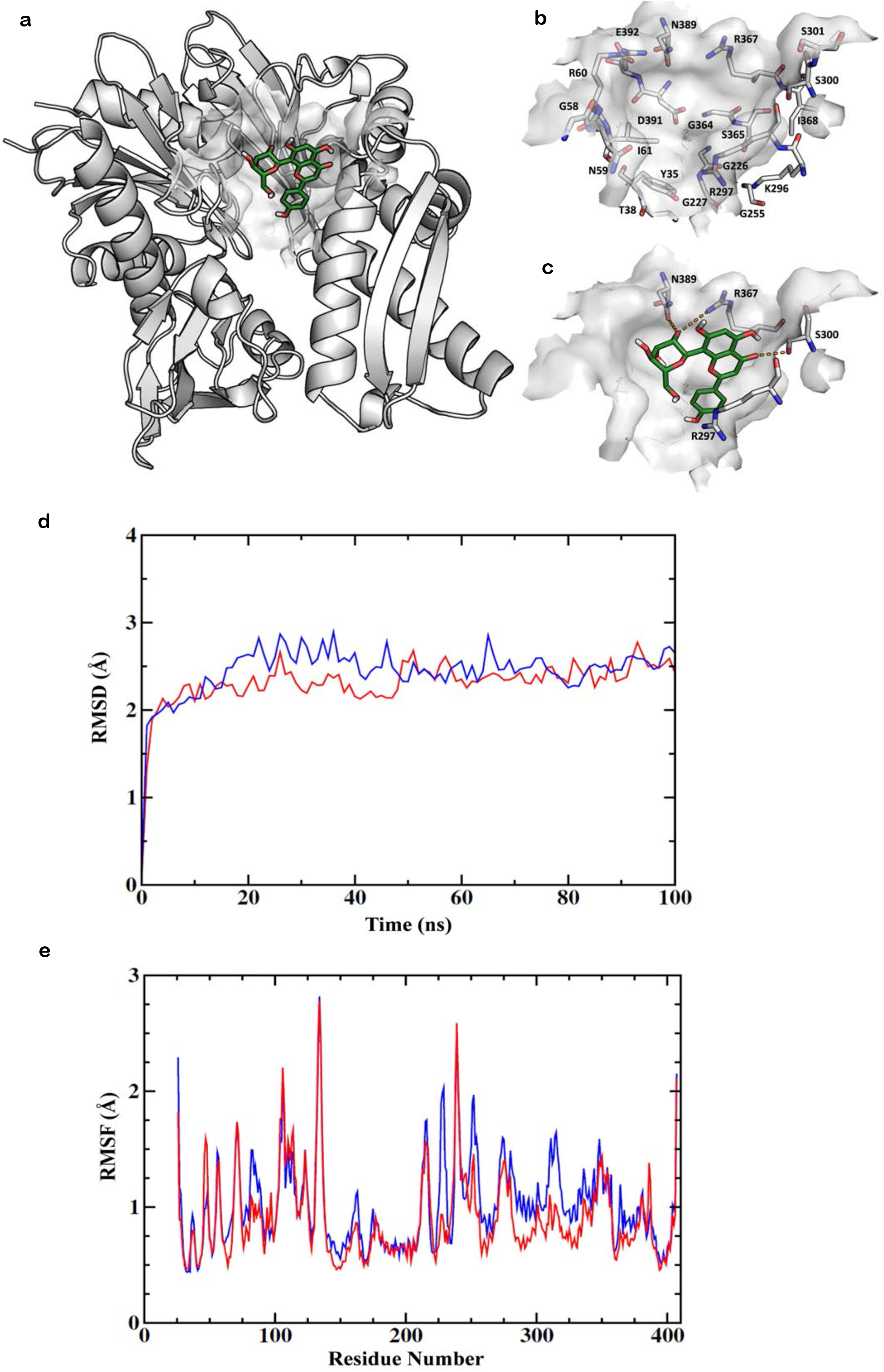
*In-Silico* molecular modeling and dynamic simulation studies identified vitexin as a potential BiP inhibitor. **(a)** Putative binding mode of the vitexin in the crystal structure of BiP. **(b)** The important residues in the binding pocket and **(c)** depicting the residues that may be important for the interaction with vitexin. Carbon atoms of vitexin are colored green and the important residues in the binding pocket are colored in gray. Oxygen atoms are colored in red, nitrogen atoms in blue, and sulfur atoms in yellow. **(d)** Root mean square deviation (RMSD) values of BiP (blue) and in complex with vitexin (red) over 100 ns. The values were obtained from the Cα atoms relative to the conformation of the first frame. **(e)** Root mean square fluctuation (RMSF) values of BiP (blue) and in complex with vitexin (red) over 100ns.

### Vitexin prevents FAM134B-BiP interaction and inhibits ER-phagy

Since our *in silico* molecular dynamics studies deciphered vitexin as a potential inhibitor of BiP, we next examined the expression of BiP in hypoxic cells treated with vitexin. Consistently, vitexin treatment downregulated BiP expression during hypoxia **(Fig 5a-5b)**. Indeed, treatment with vitexin also downregulated CI-hypoxia induced UPR at the mRNA **(Ext Fig 5a-5d)** and protein levels **(Ext Fig 5e)**. Next, we asked if treatment with vitexin prevented the interaction of FAM134B with BiP during hypoxia. Strikingly, BiP did not coimmunoprecipitate with FAM134B upon vitexin treatment in hypoxic cells **(Fig 5c)**. Furthermore, immunoblot analysis revealed that vitexin prevents autophagosomal degradation of FAM134B during CI-hypoxia **(Fig 5d and 5f)** and HE-hypoxia **(Fig 5e and 5g)**. Consistently, time-lapse imaging of vitexin-treated cells expressing GFP-WIPI-1 and mCherry-ER-3 exhibited accumulation of autophagosomes. However, autophagic flux and turnover were inhibited leading to cell death **(representative frames of time lapse imaging at different time points Ext Fig 5f and movies of vitexin treated control cells and CI-hypoxia cells are shown in supplementary videos 8 & 9 respectively)**. Collectively, these data confirm that vitexin blocks ER-phagy by inhibiting BiP from forming a complex with FAM134B.

**Fig 5:**
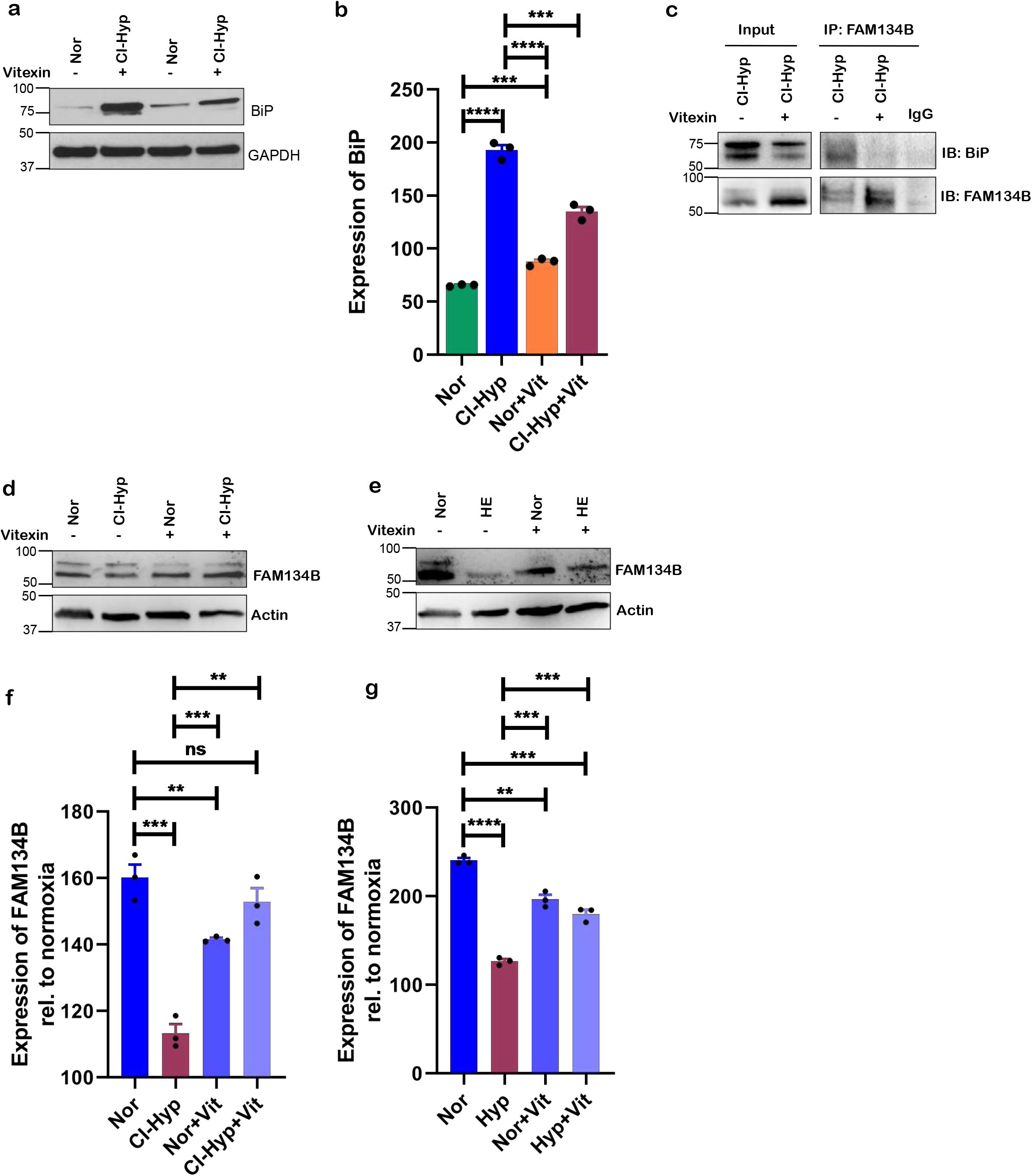
Vitexin prevents FAM134B-BiP interaction and inhibits ER-phagy. **(a)** Immunoblot of BiP upon vitexin (20µM) treatment during CI-hypoxia when compared to normoxia. **(b)** Densitometric quantification of BiP upon vitexin (20µM) treatment during CI-hypoxia when compared to normoxia. **(c)** Coimmunoprecipitation of FAM134B with BiP upon vitexin treatment during CI-hypoxia *(n=2)*. **(d)** Immunoblot of FAM134B upon vitexin treatment during CI-hypoxia and (**e)** HE (1% O2) when compared to normoxia. **(f)**Densitometric quantification of FAM134B upon vitexin treatment during CI-hypoxia and (**g)** HE (1% O2) when compared to normoxia.

### Vitexin reduces cancer cell proliferation and tumor burden in breast cancer xenograft mouse model

Having shown that vitexin inhibits ER-phagy we surmised that it could inhibit cancer cell proliferation under hypoxic stress. As expected, vitexin treatment effectively prevented the increase in cell proliferation upon hypoxic stimuli **(Fig 6a)**. As we propose that ER-phagy resolves ER-stress, we also explored if vitexin can synergistically inhibit cancer cell growth with an ER-stress inducer tunicamycin. We observed that vitexin and tunicamycin synergistically inhibited cancer cell growth **(Fig 6b)**. We next asked if vitexin can reduce tumor burden in female balb/c athymic (nuÞ/nuÞ) mice xenografted with MCF7 cells. We found that after 13, 17 and 21 days of vitexin treatment, tumor volume was significantly reduced compared to the vehicle treated mice **(Fig 6c-6d)**. Taken together, we could conclude that vitexin shows a higher therapeutic potential in treating breast cancer.

**Fig 6:**
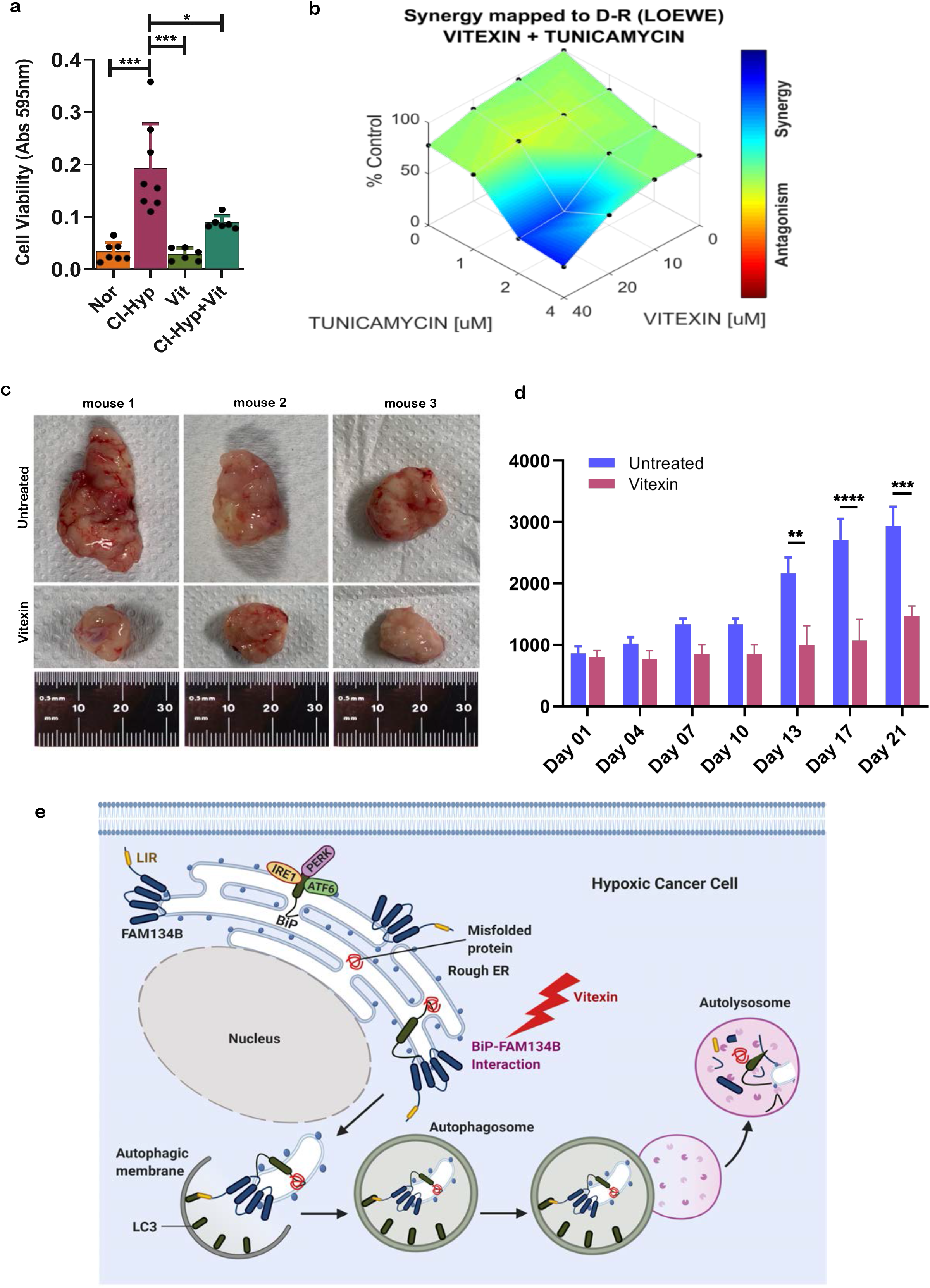
Vitexin reduces tumor burden in breast cancer xenograft mouse model. **(a)** Crystal violet cell viability assessment upon vitexin treatment during CI-hypoxia *(n=3)*. **(b)**Synergistic effect of vitexin with tunicamycin anlyzed using Combenefit® software. **(c)** Representative images of tumor grafts from female balb/c athymic (nuÞ/nuÞ) mice 21 days following an injection of vitexin and control *(n=6)*. **(d)** Tumor volumes in control and vitexin treated xenograft mice recorded on the days shown in the graph *(n=6)*. **(e)** Representation of FAM134B-BiP complex mediated ER-phagy activated upon hypoxia induced accumulation of misfolded/unfolded proteins.

## Discussion

Identification of receptors that specifically target damaged organelles and proteins into autophagosomes for degradation assists in discriminating from functional organelles. More recently, specific receptors have been identified to target damaged or excess parts of ER into autophagosomes for elimination and this process has been termed as ER-phagy. Thus far, this process has not been directly implicated in disease pathologies. Here, we report that cancer cells undergo ER-phagy regulated by FAM134B-BiP complex when they are subjected to hypoxic stress which helps the cells to mitigate ER-stress and promote cell proliferation **(Fig 6e)**.

Numerous studies have confirmed that the TME promotes cancer progression due to the ability of tumor cells to overcome stress and survive in the hostile microenvironment. Hallmark of TME is diminished levels of oxygen (less than 2%) which is essential for a cell to meet its bioenergetic needs^42^. Hypoxic conditions can be mimicked in the laboratory by either growing the cells in a hypoxic incubator with reduced levels of oxygen or treating cells with CoCl_2_^43^. Cells undergoing hypoxic stress stabilize HIF-1a which we could observe in cells subjected to CoCl_2_ treatment and those grown in a reduced Oxygen environment. Oxygen is not only required to meet the metabolic needs of the cells but is also essential for protein disulphide bond formation during protein-folding in the ER^8^ but, hypoxia disrupts protein folding resulting in the accumulation of unfolded/misfolded proteins in the ER causing ER-stress and cell death^44^. However, cancer cells alleviate ER-stress by activating UPR characterized by the expression of BiP, XBP1-s, CHOP and ATF4^45, 46^. Our data also suggest that UPR could be regulated by HIF-1a as depletion of HIF-1a abrogates the expression of UPR proteins while, the molecular mechanism by which HIF-1a regulates UPR remains to be investigated.

UPR has been shown to induce autophagy which helps in the removal of misfolded proteins and damaged organelles thus preventing DNA damage and cancer progression. On the contrary, when the cancer cells undergo nutrient starvation or hypoxia, autophagy acts as a quality control mechanism to overcome the stress and survive^19^. Consistently, autophagy is increased when cancer cells are subjected to hypoxic stress as determined by LC3I to II conversion. Autophagy encompasses autophagosome formation marked by LC3 conversion and degradation of the cargo by fusion with the lysosomes. During general/macroautophagy, LIR (LC3-interacting region) motif containing adaptor proteins such as p62 recognize ubiquitinated substrates and target them for autophagic degradation by binding to LC3^47, 48^. Upon hypoxic stress although we observed a constant turnover of WIPI which localizes to LC3 positive autophagosomes^49^, the adaptor protein p62 was not degraded. Mounting evidences suggest that autophagy selectively degrades damaged organelles to maintain cellular homeostasis. In line with this, hypoxia has been shown to target mitochondria (mitophagy)^50^ and peroxisomes (pexophagy)^51^ for autophagic degradation.

As hypoxia causes damage to ER, a failure to restore ER structure and homeostasis could be lethal to cells. Restoration of ER is achieved by the removal of excess and damaged ER caused by ER-stress. Recent evidences suggest that ER can be selectively targeted for lysosomal degradation by ER-phagy^52^. UPR has also been linked to the activation of ER-phagy however, no direct evidences linking hypoxia and ER-phagy has been reported thus far. Our data convincingly show that ER is selectively targeted for degradation in autophagosomes during hypoxia. Seven proteins containing LIR motif have been shown to function as ER-phagy receptors: CCPG1^29^, FAM134B^26^, RTN3^27^, SEC62^28^, ATL3^30^, TEX264^31, 32, 33^ and CALCOCO1^34^. These receptors have been shown to recruit subregions of the ER into autophagosomes. For instance, FAM134B is localized on the edges of the ER sheets where protein synthesis and folding occurs and has a clear role in maintaining ER volume and proteostasis^26, 27^. It is believed that the reticulon homology domain (RHD) of FAM134B aids in the fragmentation of ER and LIR motif sequesters the fragmented ER into autophagosomes during basal and starvation induced ER-phagy^25^. Hypoxia resulted in the colocalization of FAM134B with autophagosomal marker LC3 and was subsequently degraded in lysosomes indicating that cancer cells remove damaged ER by activating FAM134B-dependent ER-phagy. Silencing of FAM134B during UPR results in cell death^26^. Consistently, hypoxia stimulated ER-stress reduces the viability of FAM134B-depleted breast cancer cells. We also observed a modest increase in Sec62 which has been shown to specifically regulate the recovery of ER^28, 53^ but was not targeted for autophagic degradation during hypoxia. Notably, hypoxia upregulates FAM134B expression in chronic myeloid leukemia (CML) cells and is correlated with pro-survival^54^. It is also speculated that its upregulation is HIF-1a dependent^55^ however, silencing of HIF-1a did not alter the relative expression of FAM134B during hypoxia compared to normoxia.

FAM134B lacks intraluminal domains hence the question that remains to be answered is, how does FAM134B detect ER-stress or physiological changes within the ER? It is likely that it co-operates with accessory proteins to detect ER stress^25^. BiP and calnexin are the two major chaperone systems in the ER lumen^56, 57^ and recently it has been reported that FAM134B cooperates with calnexin which possesses a luminal chaperone domain to sense and remove misfolded procollagen^58^. BiP is a chaperone which senses accumulation of misfolded proteins in the ER resulting in the release of BiP from the UPR proteins and also chaperones the folding of accumulated proteins^13^. Strikingly, BiP strongly immunoprecipitated with FAM134B in hypoxic cells and colocalizes in the breast cancer tissues. Additionally, depletion of BiP prevented FAM134B-dependent ER-phagy and also stalled the proliferation of cancer cells subjected to hypoxia.

Though ER-phagy has been speculated to be involved in various pathologies including cancer^35^, therapeutic options to target ER-phagy has not yet been exploited. Having found that depletion of BiP prevents ER-phagy and cancer cell proliferation, we explored to pharmacologically target BiP-dependent ER-phagy. *In-silico* analysis identified vitexin, a plant derived flavone C-glycoside (apigenin-8-C-β-d-glucopyranoside)^59, 60, 61^ as an inhibitor of BiP. Although several studies have reported the therapeutic potential of vitexin in treating various medical disorders including cancer, the precise molecular target remains unresolved. We show that vitexin not only abrogates BiP dependent UPR but also inhibits FAM134B-dependent ER-phagy assisted by BiP. Decrease in UPR downstream of IRE-1 and PERK in vitexin treated cells could suggest that ER-stress is mitigated by vitexin. However, vitexin also prevents the proliferation of cancer cells which indicates that it abrogates the ability of cancer cells to overcome ER-stress. It also suggests that vitexin not only prevents BiP from binding to FAM134B but also prevents the ability of BiP to chaperone unfolded/misfolded proteins. Furthermore, induction of ER-stress with tunicamycin combined with ER-phagy inhibition using vitexin synergistically stunted cell proliferation. Therefore, we surmise that unresolved ER-stress is detrimental to cell survival and proliferation.

Disruption of ER and ER homeostasis could lead to the death of cells and cause disease pathologies. However, during the late stages of cancer when cancer cells are under various metabolic stresses including hypoxia, they adapt mechanisms such as ER-selective autophagy to overcome the damage to ER and the associated cellular processes. Therefore, targeting such adaptive mechanisms is a potential way forward to treat cancer. Our data reported here unveils FAM134B-BiP complex-mediated ER-phagy as a novel mechanism by which cancer cells prevail over hypoxia induced proteotoxic stress and targeting ER-phagy machinery as a prospective therapeutic strategy to treat cancer.

## Materials and Methods

### Cell culture

MCF-7 cells were cultured in Dulbecco’s Modified Eagle Medium (DMEM), C32 cells and U251 cells were cultured in RPMI medium supplemented with 10% fetal bovine serum and incubated at 37°C, 5% CO_2_. For hypoxia experiments, cells were incubated in the hypoxia incubator at low oxygen levels i.e. 1% O_2_.

### Drugs and treatments

CoCl_2_ 0.1M readymade solution (Cat No. 15862) and vitexin (CAS No. 3681-93-4) were procured from Sigma-Aldrich. CoCl_2_ and vitexin were used at a concentration of 500µM and 20µM respectively. Concanamycin A was purchased from Sigma Aldrich and used at a concentration of 100nM.

### Immunoblotting

MCF-7 cells were lysed in radioimmunoprecipitation assay (RIPA) buffer supplemented with protease and phosphatase inhibitors. Protein concentrations were estimated using Pierce BCA Protein assay kit (Thermo Fisher Scientific), as per the instructions. Equal amounts of proteins were separated on either 10% SDS/PAGE gels or 4-20% Mini-PROTEAN TGX Stain-Free Gels (#4568094, Bio-rad). Proteins were then transferred onto PVDF membranes and probed with the following antibodies: HIF-1α (D2U3T) (#14179, Cell Signaling Technology), BiP (C50B12) (#3177, Cell Signaling technology), CREB-2 (SC-200, Santacruz), CHOP (#2895, Cell Signaling Technology), LC3B (#83506, Cell Signaling Technology), ER Stress Antibody Sampler Kit (#9956, Cell Signaling Technology), Sec24C (#14676, Cell Signaling Technology), Lamin B1 (#12586, Cell Signaling Technology), CCPG1 (ab150465, Abcam), Sec62 (ab137022, Abcam), FAM134B (#61011, Cell Signaling Technology) and anti-FAM134B polyclonal antibody (a kind gift from Ivan Dikic), RTN3 (# PA578316 Thermo Fisher Scientific), Normal Rabbit IgG (#2729, Cell Signaling Technology), XBP-1s (#12782, Cell Signaling Technology), SQSTM1/p62 (#5114, Cell Signaling Technology). Beta actin, calnexin (#2679, Cell Signaling Technology) or GAPDH (sc-32233, Santacruz) were used as loading controls. After incubation with secondary horseradish peroxidase (HRP)-conjugated antibodies, the blots were washed and developed using enhanced chemiluminescence reagent in the Chemidoc MP or ImageQuant LAS4000.

### Immunofluorescence staining and confocal microscopy

MCF-7 cells grown on the glass coverslips were treated with CoCl2 and vitexin for 16-24 h and fixed with 4% formaldehyde in PBS for 15 min at RT. The cells were then permeabilized with 0.3% Triton X-100 in PBS for 5 min and blocked with 3% BSA for 1 h at RT. The cells were incubated overnight with primary antibody against HIF-1a, BiP, CREB-2, CHOP, LC3B, calnexin and FAM134B at 4°C. After overnight incubation, the cells were washed with PBS and incubated with either Alexa Fluor 594-conjugated goat anti-rabbit/anti-mouse or Alexa Fluor 488-conjugated goat anti-rabbit/anti-mouse secondary antibody for 1h at RT in the dark. The cells were then washed, and coverslips were mounted using ProLong Diamond antifade containing DAPI to stain the nuclei. Staining of mouse and human breast cancer tissues was performed after antigen retrieval using 0.01M Citrate buffer pH 6.0. The slides were imaged under Leica SP8 confocal or Leica THUNDER imager. Human breast cancer tissues were obtained with consent from the patients which was approved by the Ethical committee of the University of Campania “Luigi Vanvitelli” (Prot. 71-13/2/2129).

### Quantitative real-time PCR

Total RNA from MCF-7 cells (1×10^6^ cells/well) was isolated using RNeasy Mini kit (74106; Qiagen), and 500ng cDNA was synthesized with random hexamers by reverse transcription (SuperScript III; 18080; Invitrogen). 20µL of PCR reactions contained10ng cDNA, 0.4 µmol/liter of each forward and reverse primer, and master mix (SsoFast EvaGreen Supermix; 1725201; Bio-Rad). Real-time PCR was performed under the following conditions: initial denaturation step at 95°C for 2 min and 40 cycles at 95°C for 5 s and 60°C for 15 s, followed by a denaturation step at95°C for 60 s and a subsequent melt curve analysis to check amplification specificity. Results were analyzed by the comparative threshold cycle method with hypoxanthine-guanine phospho ribosyl transferase (*HPRT*) as the endogenous reference gene for all reactions. The relative mRNA levels of untreated samples were used as normalized controls for the CoCl2 and vitexin treated samples. All reactions were performed in triplicate and a non-template control was included in all experiments to exclude DNA contamination. Primer sequences are listed in Table 1.

### Time-Lapse imaging for cell proliferation

MCF-7 cells were seeded (2×10^4^ cells/well) in the ibidi µ slide 8 well (cat no.80827) suitable for live cell imaging in the CellVoyager CV1000 confocal imaging system (Yokogawa). A day after, the cells were treated with appropriate concentrations of CoCl_2_ and vitexin and set up for time-lapse imaging over 24 h duration at an interval of 20min.

### Time-Lapse Imaging for ER-phagy

MCF-7 cells were transfected with mCherry-ER-3 and GFP-WIPI-1 plasmids and seeded (2×10^4^ cells/well) in the Nunc™ Lab-Tek™ II Chamber Slide (cat no. 154526) suitable for live imaging. The next day, following treatment with CoCl2 and vitexin cells were set up for time-lapse imaging in the Leica SP8 confocal or Leica THUNDER imager over 24 h duration at an interval of 20min. The data was processed and analyzed using imageJ software.

### siRNA and Plasmid transfection experiments

MCF-7 cells (0.5×10^6^ cells/well) were incubated with either 100nM nontargeting siRNA (SR-CL000-005; Silencer™ Select) or 100nM siRNA specific for HIF-1a (HIF1a; L-004018-00-0005; Dharmacon), BiP (siRNA ID: s29012; Silencer™ Select)and FAM134B (siRNA ID: s29012, s29013; Silencer™ Select) together with the transfection reagent Lipofectamine 3000 (L3000-008; Invitrogen) for 48 h according to the manufacturer’s instructions. Knockdown efficiency was assessed by Western blot analysis using antibodies against HIF-1a, BiP and FAM134B respectively. mCherry-ER-3was a gift from Michael Davidson (Addgene plasmid # 55041; http://n2t.net/addgene:55041; RRID:Addgene_55041) andpMXs-IPGFP-WIPI-1^62^was a gift from Noboru Mizushima (Addgene plasmid # 38272; http://n2t.net/addgene:38272; RRID:Addgene_38272) plasmids were transiently transfected into the MCF-7 cells for 48h using Lipofectamine 3000.

### Immunoprecipitation

MCF-7 cells (5.0 × 10^6^ cells/well) were lysed with RIPA buffer containing protease and phosphatase inhibitors. After preclearing the cell lysate with protein A/G agarose magnetic beads (16-663; Millipore) for 1 h, beads were removed by placing the tube on a magnetic rack. The whole cell lysate (∼1000 µg of protein) was incubated overnight at 4°Cwith 4 µg of an antibody against FAM134B. Protein A/G agarose beads were added again and incubated for an additional 1 h at room temperature. The immunoprecipitated proteins along with the agarose beads were collected by placing the tube on a magnetic rack. The collected beads were washed three times with RIPA buffer. The washed samples were mixed with SDS-PAGE sample loading buffer, boiled, and resolved on a 10% SDS–polyacrylamide gel. The respective proteins precipitated were probed for specific antibodies for immunoblot analysis.

### Crystal Violet cell viability assay

MCF-7cells were cultured in 96-well plate at a density of 1×10^4^ cells/well. Some wells were kept without cells to serve as control for non-specific binding of the crystal violet. After 16-24 h, medium was aspirated and added 100µL of fresh medium supplemented with appropriate concentrations of drugs (CoCl_2_ and vitexin) and incubated for 24 h at 37°C in standard culture conditions. After 24 h of incubation, wells were gently washed twice with water and incubated with 50µL 0.5% crystal violet staining solution for 20min at room temperature on a bench rocker at a frequency of 20 oscillations per minute. Then the plate was air-dried without the lid for 2 h at room temperature and incubated with 200µL methanol for 20min at room temperature on a bench rocker at a frequency of 20 oscillations per minute. Absorbance was measured at 570nm using a microplate reader (Bio Tek™ EPOCH).

### *In-Silico* molecular docking studies

#### Ligand preparation

The 2D structures of the selected ligands were drawn using ChemSketch software (https://chemsketch.en.softonic.com/). The ligands were prepared using Ligprep of Schrödinger suite. Bond orders were refined, missing hydrogen atoms were added followed by generating 3D structures with possible ligand ionization and tautomeric states at pH 7.0 ± 2.0 using Epik module. The generated low energy conformers were finally energy minimized by using OPLS_2005 force field.

#### Protein preparation for *in-silico* docking studies

The 3D X-ray structure of human GRP78 ATPase domain complexed with 2’-deoxy-ADP and inorganic phosphate (5F0X.*pdb*, Resolution: 1.6 Å) was retrieved from protein data bank and was further prepared using protein preparation wizard of Schrödinger suite 2015-3. The initial protein structure was a homo dimer, where the redundant chains have been removed with deleting waters, refining bond orders and addition of hydrogens. Prime module was used for adding missing side chains and loops followed by generating protonation and tautomeric states of acidic and basic residues at normal pH 7.0 by PROPKA. Next, protein hydrogen bond assignment was done along with side chain flipping of His, Asp and Glu with reorienting hydroxyl and thiol groups. Finally, protein minimization was performed using OPLS_2005 (Optimized Potentials for Liquid Simulations) molecular force field with RMSD of crystallographic heavy atoms kept at 0.30 Å. The quality of prepared protein was validated using Ramachandran plot.

### Grid generation and molecular docking

A Grid box was generated at the centroid of active site keeping receptor van der Waals scaling of 1.0 with partial charge cutoff at 0.25. The generated low energy conformers were docked into the active site of 5F0X.*pdb* using extra precision mode (XP) docking of Glide (Glide v 6.8, Schrödinger 2015-3) keeping default parameters. The docked pose was selected based on terms of Glide g score, Glide model and Glide energy values.

### Cell viability assay and synergism with tunicamycin

Cell viability was measured by the colorimetric 3-(4,5-dimethyl-2-thiazolyl)-2,5-diphenyltetrazolium bromide (MTT) assay. Cells were seeded in 96-well plates at a density of 10^4^ cells per well and treated with vitexin and tunicamycin. 100 μL of 1 mg/mL MTT (Sigma) in DMEM medium containing 10% fetal bovine serum was added to treated cells for 4 h at 37°C. The medium was replaced with 200μL of DMSO and shaken for 15 min, then absorbance at 540 nm was measured using a microplate ELISA reader with DMSO used as the blank. To quantify the synergistic or antagonist effect of the drugs combinations, Combenefit® software was used.

### In vivo mouse xenograft and vitexin treatment

Four-to six-week-old female balb/c athymic (nuÞ/nuÞ) mice were purchased from The Charles River Laboratories. The research protocol was approved, and mice were maintained in accordance with the institutional guidelines of the Università degli Studi della Campania L. Vanvitelli Animal Care and Use Committee. Animal care was in compliance with Italian (Decree 116/92) and European Community (E.C. L358/1 18/12/86) guidelines on the use and protection of laboratory animals. Mice were acclimatized at Università degli Studi della Campania L. Vanvitelli Medical School Animal Facility for 1 week prior to being injected with cancer cells and then caged in groups of 3. A total of 5× 10^6^ MCF-7 cells were resuspended in 200 μL of Matrigel (BD Biosciences) and PBS (1:1) and implanted subcutaneously into the right flank of 12 nude female mice. At week 2, once tumors reached a mean volume of 600 mm^3^, mice were randomized into treatment group (6 mice) or control group (6 mice), to receive treatment with vitexin 2 mg/kg or vehicle (dimethyl sulfoxide (DMSO)), respectively, via intraperitoneal injection, 5 days a week, for 3 weeks. Tumor size was evaluated twice a week by caliper measurements using the following formula: π/6 x larger diameter x (smaller diameter)^2^. Tumor response was assessed by using volume measurements and adapted clinical criteria.

## Statistical Analysis

Statistical analyses were performed using GraphPad Prism Software (Version 8.0). Unpaired Student’s *t* test or two-way anova with Bonferroni Posthoc test was conducted for all the datasets as indicated in figure legends to determine statistical significance. All the data are represented as mean± SEM. For all tests, a *P* value <0.05 was considered statistically significant (* p<0.05; ** p<0.01; *** p<0.001; **** p<0.0001).

## Acknowledgements

We thank the Robinson laboratory members and Dr Alexandra Stolz, Institute of Biochemistry 2, Goethe University School of Medicine, Frankfurt, Germany for helpful discussions and critical review of the manuscript; Dr Makoto Kamei, Imaging Facility, Centre for Cancer Biology, Adelaide. The imaging facility is supported by funding from Australian Cancer Research Foundation. Dr Chia Chi, Centre for Cancer Biology for helping us to set up hypoxic conditions and Dr Srikanth Jupudi, JSS College of Pharmacy for his help with the In-silico docking studies. Work in the laboratory of NR is supported by funds from Centre for Cancer Biology and University of South Australia. Work in the laboratory of NR was also supported by funds from Cologne Excellence Cluster on Cellular Stress Responses in Aging-Associated Diseases (CECAD; funded by the DFG within the Excellence Initiative by the German federal and state governments) and Köln Fortune, University clinic, Cologne, grants from Deutsche Forschungsgemeinschaft (SFB 670). The authors acknowledge Department of Science and Technology, Government of India for financial support vide reference no SR/WOS-A/LS-21/2016 under Women Scientist Scheme to carry out research in this area to SC. SC was also supported with a short-term travel fellowship from European Association for Cancer Research.

